# Separated and overlapping neural coding of face and body identity

**DOI:** 10.1101/2021.01.30.428936

**Authors:** Celia Foster, Mintao Zhao, Timo Bolkart, Michael J. Black, Andreas Bartels, Isabelle Bülthoff

## Abstract

Recognising a person’s identity often relies on face and body information, and is tolerant to changes in low-level visual input (e.g. viewpoint changes). Previous studies have suggested that face identity is disentangled from low-level visual input in the anterior face-responsive regions. It remains unclear which regions disentangle body identity from variations in viewpoint, and whether face and body identity are encoded separately or combined into a coherent person identity representation. We trained participants to recognize three identities, and then recorded their brain activity using fMRI while they viewed face and body images of the three identities from different viewpoints. Participants’ task was to respond to either the stimulus identity or viewpoint. We found consistent decoding of body identity across viewpoint in the fusiform body area, right anterior temporal cortex, middle frontal gyrus and right insula. This finding demonstrates a similar function of fusiform and anterior temporal cortex for bodies as has previously been shown for faces, suggesting these regions may play a general role in extracting high-level identity information. Moreover, we could decode identity across neural activity evoked by faces and bodies in the early visual cortex, right inferior occipital cortex, right parahippocampal cortex and right superior parietal cortex, revealing a distributed network that encodes person identity abstractly. Lastly, identity decoding was consistently better when participants attended to identity, indicating that attention to identity enhances its neural representation. These results offer new insights into how the brain develops an abstract neural coding of person identity, shared by faces and bodies.

## 1. Introduction

Being able to recognise the identity of people we encounter in our daily life is a crucial ability for our social interactions. We use multiple sources of information in order to encode and then subsequently recognise specific people, including facial information, body information, and face and body movements (Dobs, Bülthoff, & Schultz, 2016; Hahn, O’Toole, & Phillips, 2015; O’Toole et al., 2011; O’Toole, Roark, & Abdi, 2002; Rice, Phillips, Natu, An, & O’Toole, 2013; Rice, Phillips, & O’Toole, 2013; Robbins & Coltheart, 2012). For instance, our visual system is remarkably good at recognising the identity of familiar people, irrespective of changes in viewpoint, illumination, position, pose and expression. This seemingly effortless ability is computationally challenging, as these changes lead to a great variability in low-level visual information arriving on the retina, yet we are able to distinguish between identities that look comparably similar to one another. Although the face gives strong cues for recognition, body information also contributes to the visual recognition of a person’s identity, especially when face information is not optimal for recognition, for example when a person is far away (Hahn et al., 2015; O’Toole et al., 2011; Rice, Phillips, Natu, et al., 2013; Rice, Phillips, & O’Toole, 2013). It is not yet fully understood how person identity is encoded in the brain, and whether the neural coding of face identity and body identity is separated or overlapping. We aimed to address these questions in the present study.

Neural coding of face identity has been associated with face-responsive brain regions in the fusiform gyrus and anterior temporal cortex (Haxby, Hoffman, & Gobbini, 2000). These regions respond when participants recognize face identities (Grill-Spector, Knouf, & Kanwisher, 2004; Hoffman & Haxby, 2000; Nasr & Tootell, 2012) and dysfunction of these regions can lead to impairments in face recognition ability (Barton, 2008; Busigny et al., 2014; Hadjikhani & de Gelder, 2002; Jonas et al., 2015). Anterior temporal cortex is thought to be of particular importance in encoding high-level face identity representations. Patterns of activity in this region can distinguish between different face identities (Kriegeskorte, Formisano, Sorger, & Goebel, 2007). Moreover, neural patterns evoked by different face identities in this region can generalise across face viewpoint (Anzellotti, Fairhall, & Caramazza, 2014; Freiwald & Tsao, 2010; Guntupalli, Wheeler, & Gobbini, 2017), face expression (Nestor, Plaut, & Behrmann, 2011) and different halves of the same face (Anzellotti & Caramazza, 2016). The fusiform face area (FFA) also responds to changes in identity (Andrews & Ewbank, 2004; Gauthier et al., 2000; Loffler, Yourganov, Wilkinson, & Wilson, 2005; Rotshtein, Henson, Treves, Driver, & Dolan, 2005; Winston, Henson, Fine-Goulden, & Dolan, 2004), and some studies have shown that face identity responses in the FFA can generalise across viewpoint (Anzellotti et al., 2014; Guntupalli et al., 2017). Other studies have also found high-level face identity responses in the occipital face area (OFA) (Anzellotti et al., 2014), the superior intraparietal sulcus (Jeong & Xu, 2016) and right inferior frontal cortex (Guntupalli et al., 2017).

Although psychological research has shown that we also use body information to recognise people (Hahn et al., 2015; O’Toole et al., 2011; Rice, Phillips, Natu, et al., 2013; Rice, Phillips, & O’Toole, 2013; Robbins & Coltheart, 2012), much less is known about the brain regions encoding body identity. An fMRI repetition suppression study found lower responses in the extrastriate and fusiform body areas (EBA and FBA) to repeated presentation of the same body identity as compared to presentation of different body identities, suggesting that these regions encode body identity information (Ewbank et al., 2011). Stronger neural responses to the bodies of familiar people, as compared to unfamiliar people, have been observed in the FBA as well as the inferior and medial frontal gyrus, cingulate gyrus, central and post-central sulcus and inferior parietal lobe (Hodzic, Kaas, Muckli, Stirn, & Singer, 2009). However, no study has investigated which regions of the human brain contain different patterns of neural activity evoked by different body identities, or furthermore which brain regions contain patterns of responses to different body identities that can generalize across different viewpoints. In macaques, electrophysiological recordings have shown that the body-responsive patches contain body identity information that can generalize across viewpoint and pose (Kumar, Popivanov, & Vogels, 2019). Interestingly, identity decoding accuracy was higher in the more anterior body patch, suggesting an important role of more anterior temporal regions in encoding viewpoint-invariant body identity, similar to the function of more anterior face-responsive regions in viewpoint-invariant coding of face identity.

In the present study, we aimed to address two fundamental questions about the neural representation of face and body identity. Firstly, we investigated which brain regions encode body identity, and which regions encode body identity in a viewpoint-invariant manner. Secondly, we investigated whether face- and body-based identity information are encoded in separated or overlapping brain networks. Of particular interest is where in the brain face and body identity information from the same person is combined into a stimulus-independent person identity representation. It has been suggested that brain regions processing faces and bodies in occipitotemporal cortex are mostly separated, parallel networks (Pitcher, Charles, Devlin, Walsh, & Duchaine, 2009; Premereur, Taubert, Janssen, Vogels, & Vanduffel, 2016; Schwarzlose, Baker, & Kanwisher, 2005). Our recent study showed that certain aspects of face and body information (e.g. weight) are integrated in the EBA (Foster et al., 2019). In macaques, the anterior face patches show stronger neural responses to images of a whole person than to the addition of the responses to the face and body shown alone (Fisher & Freiwald, 2015), suggesting that these regions may integrate face and body information. If face and body information is integrated to form an abstract neural person identity code, we would expect similar patterns of neural responses to a particular identity, regardless of whether the person is viewed from an image of their face or body.

To address these questions, we trained participants to recognize three identities and then recorded their brain activity using fMRI as they viewed images of the face and body of these three identities from three different viewpoints. Participants performed two behavioural tasks during the experiment, one where they responded to the stimulus identity (i.e. identity recognition task) and the other where they responded to the stimulus viewpoint (i.e. viewpoint recognition task). This manipulation allowed us to investigate if neural coding of person identity is enhanced when participants attend to identity as compared to when they do not (i.e. when they attend to viewpoint). First, to investigate which brain regions contained patterns of neural activity that could distinguish between face identities and between body identities, we trained linear support vector machine (SVM) classifiers to distinguish between patterns of activity evoked by the identities, and then tested these classifiers on their ability to decode face and body identities in a separate test set of data. Second, to test which brain regions contain patterns of brain activity that encode face or body identities in a viewpoint-invariant manner, we trained classifiers using neural activity evoked by the face/body identities from two viewpoints and then tested their ability to distinguish between neural activity evoked by the face/body identities from the third viewpoint (i.e. identity classifiers that could generalize across viewpoint). Third, to test which brain regions encoded person identity using an abstract code, independent of the stimulus type (i.e. faces or bodies), we trained a classifier using neural activity evoked by the face identities and then tested this classifier on its ability to distinguish between neural activity evoked by the body identities, and vice versa. We performed all of these analyses in face- and body-responsive regions of interest (ROIs) as well as in whole-brain searchlight analyses, and we performed these analyses separately on fMRI data where participants performed the identity and viewpoint recognition tasks.

## 2. Materials and methods

### 2.1. Participants

Twenty participants completed the experiment. One participant was excluded from the data analyses due to poor performance in the behavioural task (< 40% correct responses in one condition). The remaining 19 participants (13 female, 6 male, 21-51 years old) were included in the behavioural and fMRI analyses presented here. The experiment procedure was approved by the local ethics committee of the University Clinic Tübingen and was conducted according to the Declaration of Helsinki. All participants provided written informed consent prior to the start of the experiment.

### 2.2. Stimuli

#### 2.2.1. Main experiment stimuli

Our stimuli (Fig. 1A) consisted of separate face and body images of three identities from three viewpoints; 0° (front), 45° (three-quarter) and 90° (profile). The three identities were all female, to ensure that sex did not differ between the three identities. For each identity, we recorded both a 3D face scan with a neutral expression and a 3D body scan in an A-pose. The face scans were then aligned to a 3D shape and expression model (Li, Bolkart, Black, Li, & Romero, 2017) and the body scans were aligned to a 3D shape and pose model (Loper, Mahmood, Romero, Pons-Moll, & Black, 2015). We then generated images of the three individuals from the three viewpoints (0°, 45° and 90°). For body images, we covered the face using a grey rectangle to remove face information from the body images.

**Figure 1.** Experimental stimuli and procedure. (A) Stimuli were face and body images of three female identities shown from three viewpoints (0° and 45° shown here). (B) Example block of stimuli shown in the fMRI experiment. Participants viewed 6 images from one condition (i.e. face or body, one identity, one viewpoint) within a block, which varied in their image size (2 repetitions of 3 image sizes, shown in a random order). Participants performed two tasks; they responded immediately when they saw an image of the smallest image size, and they responded at the end of the block during fixation to indicate which identity or viewpoint was shown in the block (half of the experiment trials were the identity recognition task, and the other half of the trials were the viewpoint recognition task).

For each identity, we also recorded a short video showing the whole body with the head fully visible turning between the left and right profile view. This video was used for identity learning prior to the fMRI experiment (see procedure below).

#### 2.2.2. Localizer stimuli

Stimuli used to localize face- and body-responsive regions of interests were grayscale images of faces, headless bodies, objects and phase-scrambled images. The phase-scrambled images were generated by creating Fourier-scrambled versions of an image consisting of a collage of the face and headless body images.

### 2.3. Experimental procedure

The study consisted of a short identity learning session (outside of the MRI scanner) followed immediately by the main fMRI session, which consisted of eight runs of the main experiment and one run of a localizer experiment.

#### 2.3.1. Identity learning

We trained participants to recognise the three identities from images of their face and body. The identity learning session consisted of five repetitions of a learning and testing with feedback procedure. During learning, participants viewed a 15 s video of each identity (showing their whole body turning between the left and right profile), then viewed the separate face and body images of the identity from the three viewpoints (0°, 45° and 90°), until the participant pressed a button to continue. A name was presented above all the images of each identity, so that participants could learn to associate each identity with its name. Following learning, participants completed 54 trials of the testing procedure with feedback. The 54 trials consisted of three repetitions of the 18 stimulus conditions (face or body, three identities, three viewpoints) presented in a random order. In each trial, participants viewed a fixation cross for 1 s, then a stimulus image for 1 s, then a grey screen. Participants had up to 6 s to respond using a button press to indicate which identity was shown. After making a response, participants were given feedback as to whether their response was correct or not. At the end of the test, participants were shown an overall percentage correct score.

The identity learning session was presented on a laptop with resolution 1366×768, running Windows 10 with Matlab 2014a using the Psychophysics Toolbox extensions (Brainard, 1997; Kleiner, Brainard, & Pelli, 2007; Pelli, 1997).

#### 2.3.2. Main fMRI experiment

Participants lay supine in the MRI scanner and viewed the stimuli on a screen positioned 92 cm behind their head, via a mirror attached to the head coil. We presented the stimuli in a block design, with each block showing images from 1 of 18 conditions of a 2 (face or body) x 3 (identity) x 3 (viewpoint) factorial design. Each run contained 3 repetitions of all 18 conditions presented in a random order. The 18 conditions were preceded by and followed by 8 s of fixation.

Each block contained 6 images varying in their image size (Fig. 1B). There were 2 repetitions of 3 image sizes presented in a random order. The three image sizes had scale factors of 1, 1.3 and 1.6 (i.e. the largest image size was 1.6 times the width and height of the smallest image size). For face stimuli the mean widths and heights of the 3 image sizes were 4.4° x 6.4°, 3.6° x 5.2° and 2.8° x 4.0° of visual angle, for body stimuli the mean widths and heights of the 3 image sizes were 3.2° x 7.7°, 2.6° x 6.2° and 2.0° x 4.8° of visual angle. Each image was shown for 900 ms and a 100 ms blank screen was shown between images. Each block was followed by 2 s fixation.

The experiment was programmed with Matlab 2017a using the Psychophysics Toolbox extensions (Brainard, 1997; Kleiner et al., 2007) on Ubuntu 17.10. The experiment was presented using a projector with resolution 1920×1080 onto a screen with a width and height of 25° x 14° of visual angle.

Participants performed an identity recognition task in half of the experiment runs and a viewpoint recognition task in the other half of the experiment runs. Each run began with an instruction informing the participant whether they should respond to the identity or the viewpoint of the stimuli in that run. Participants responded during fixation at the end of each block by pressing a corresponding button to indicate which identity (ID1, ID2 or ID3) or which viewpoint (0°, 45° or 90°) was shown in the block. To ensure participants kept their attention on the stimuli throughout each block, we instructed participants to immediately press a button with their thumb whenever they saw an image that was shown in the smallest of the three image sizes.

#### 2.3.3. fMRI localizer experiment

Participants completed one run of a localizer experiment which was used to define face- and body-responsive brain regions. Participants viewed face, body, object and phase-scrambled images in a block design. Each block consisted of 8 images, which were each shown for 1.8s followed by a 0.2 s blank screen. Blocks were presented in a carryover counterbalanced sequence, such that face, body, object and phase scrambled blocks were preceded by each other block type an equal number of times (Brooks, 2012). Face, body and object images were shown in front of the phase-scrambled images to keep the area of retinal stimulation the same for all blocks. Participants performed a one-back matching task on the images to keep their attention on the stimuli. Images were repeated on average once every 9 s.

### 2.4. MRI sequence parameters

MRI data was acquired with a 3T Siemens Prisma scanner and a 64-channel head coil (Siemens, Erlangen, Germany). Functional T2* echoplanar images (EPI) were acquired using the following sequence parameters; multiband acceleration factor 2, GRAPPA acceleration factor 2, TR 1.84 s, TE 30 ms, flip angle 79°, FOV 192×192 mm. Volumes consisted of 60 slices and had an isotropic voxel size of 2×2×2 mm. We discarded the first 8 volumes of each run to allow for equilibration of the T1 signal. We additionally acquired a high-resolution T1-weighted anatomical scan for each participant with the following sequence parameters; TR 2 s, TE 3.06 ms, FOV 232×256 mm, 192 slices, isotropic voxel size of 1×1×1 mm.

### 2.5. MRI data preprocessing

We preprocessed our MRI data using SPM12 (http://www.fil.ion.ucl.ac.uk/spm/). Functional images were slice-time corrected, realigned and coregistered to the anatomical image. Functional images from the localizer experiment were additionally smoothed with a 6 mm Gaussian kernel. ROI and searchlight analyses on functional images from the main experiment were conducted on unsmoothed data in subject-space. The resulting searchlight classification accuracy maps were then normalised to MNI (Montreal Neurological Institute) space, and spatially smoothed with a 6 mm Gaussian kernel. For the whole-brain univariate analyses the coregistered data was normalized to MNI space and spatially smoothed with a 6 mm Gaussian kernel.

### 2.6. Definition of regions of interest

Using fMRI data from the localizer experiment, we defined three face-responsive ROIs (the OFA, FFA and ATFA) and two body-responsive ROIs (the EBA and FBA), see Table 1. We first attempted to define the face-responsive ROIs using the contrast faces > objects and the body-responsive ROIs using the contrast bodies > objects. If we could not define a ROI in a participant using this contrast, we then attempted to define the ROI using the contrast faces > scrambled images or bodies > scrambled images. We initially used a contrast threshold of *p* < .001 (uncorrected) and reduced the threshold to *p* < .01 (uncorrected) if the ROI could not be defined with the initial threshold.

**Table 1.** Mean MNI coordinates and volume of each ROI, ± standard deviations. N shows the number of participants each ROI was identified in.

| ROI | hem | x |  | y |  | z |  | Volume (mm<sup>3</sup>) | N |
| --- | --- | --- | --- | --- | --- | --- | --- | --- | --- |
| OFA | left | -35 | ± 6.9 | -86 | ± 5.9 | -11 | ± 3.6 | 731 ± 346.5 | 19 |
| | right | 38 | ± 4.1 | -81 | ± 6.0 | -10 | ± 3.3 | 994 ± 382.8 | 19 |
| FFA | left | -40 | ± 2.8 | -55 | ± 5.5 | -20 | ± 2.8 | 709 ± 364.3 | 19 |
| | right | 42 | ± 3.3 | -52 | ± 4.3 | -18 | ± 2.4 | 1083 ± 400.9 | 19 |
| ATFA | left | -34 | ± 5.5 | -11 | ± 6.7 | -33 | ± 6.9 | 177 ± 120.6 | 14 |
| | right | 34 | ± 5.8 | -8 | ± 5.5 | -37 | ± 5.8 | 335 ± 265.6 | 18 |
| EBA | left | -44 | ± 3.7 | -78 | ± 5.4 | 3 | ± 6.6 | 896 ± 486.0 | 19 |
| | right | 49 | ± 2.3 | -70 | ± 2.7 | 0 | ± 4.7 | 1686 ± 453.0 | 19 |
| FBA | left | -39 | ± 4.2 | -50 | ± 6.5 | -20 | ± 3.0 | 703 ± 459.0 | 18 |
| | right | 40 | ± 3.9 | -50 | ± 5.6 | -19 | ± 2.4 | 1148 ± 552.6 | 19 |

### 2.7. Behavioural analyses

We calculated participants’ accuracy in the identity and viewpoint recognition tasks using % correct. To investigate if stimulus identity affected participants’ ability to recognize the identity or viewpoint of the stimuli during the fMRI experiment, we performed one-way repeated-measures ANOVAs with three levels (ID1, ID2 and ID3), separately for identity and viewpoint recognition of the face and body stimuli. Prior to each ANOVA, we tested for non-sphericity using a Mauchly’s test of sphericity, and where necessary we corrected for non-sphericity using a Greenhouse-Geisser correction.

### 2.8. Univariate fMRI analyses

We conducted univariate analyses to investigate if there were any differences in the mean BOLD signal evoked by the three stimulus identities. To do this, we used SPM12 to model the fMRI data with a GLM. The GLM contained regressors for each of the experimental conditions. We then performed one-way repeated-measures ANOVAs with three levels (ID1, ID2 and ID3) separately for face and body stimuli, in face- and body-responsive ROIs and in whole-brain analyses. For ROI analyses, we tested for non-sphericity using a Mauchly’s test of sphericity, and where necessary corrected for non-sphericity using a Greenhouse-Geisser correction. We then assessed significance using a threshold of *p* < .05, Bonferroni-corrected for N = 5 ROIs. Following any significant ANOVA results, we performed follow-up paired *t*-tests between the three identities to determine between which identities there were differences in neural activation. For whole-brain analyses, we assessed significance using a threshold of *p* < .05, false discovery rate (FDR) corrected.

### 2.9. Multivoxel pattern analyses (MVPA)

We conducted multivoxel pattern analyses (MVPA) to investigate if there were differences in the patterns of neural activity evoked by the three stimulus identities. To do this, we first used SPM12 to model the fMRI data with a GLM. This GLM contained one regressor for each stimulus block. We then performed MVPA analyses on the beta weight images from the GLM using The Decoding Toolbox (Hebart, Görgen, & Haynes, 2015). We feature-scaled the data using z-score normalisation, where we estimated the mean and standard deviation on the training data and applied these values to the test data. Any outlier values (greater than 2 standard deviations from the mean) were set to 2 or −2. We performed 3 different classification analyses using a linear SVM classifier (LIBSVM).

We performed all classification analyses in face- and body-responsive brain regions and in whole-brain searchlight analyses (4-voxel radius). For ROI analyses, significance was determined using permutation testing. Each analysis was repeated 10,000 times with the condition labels randomly assigned to generate a null distribution of mean classification accuracies expected by chance. We assessed significance by comparing how often we obtained a mean classification accuracy in the null distribution greater than or equal to the actual mean classification accuracy obtained for that ROI. We assessed significance using a threshold of *p* < .05, and used a Bonferroni-correction for N = 5 ROIs tested.

For searchlight analyses, we performed group analyses using nonparametric permutation tests with SnPM13 (http://warwick.ac.uk/snpm). We performed 10,000 permutations for each analysis and used 6 mm FWHM variance smoothing. We assessed significance with a threshold of *p* < .05, FDR corrected.

#### 2.9.1. Identity classification analyses

We performed identity classification analyses to investigate which brain regions contain different patterns of activity evoked by different identities. We performed these analyses separately for neural activity evoked by face and body stimuli, and when participants performed the identity and viewpoint recognition tasks. We trained a linear SVM classifier to distinguish between patterns of neural activity evoked by the three identities using three runs of fMRI data. We then tested the classifier on its ability to predict the stimulus identities from neural activity in the fourth run of data. We performed a four-fold cross-validation procedure (where each run was used as the held-out test dataset once), and we determined the final decoding accuracy by averaging over the four cross-validation iterations.

#### 2.9.2. Viewpoint-invariant identity classification analyses

We performed viewpoint-invariant identity classification analyses to investigate which brain regions contain patterns of neural activity evoked by the stimulus identities that can generalize across stimulus viewpoint. As previously, we performed these analyses separately for neural activity evoked by face and body stimuli, and when participants performed the identity and viewpoint recognition tasks. In these viewpoint-invariant analyses, we used three runs of fMRI data to train a linear SVM classifier to distinguish between patterns of neural activity evoked by the three identities from two of three viewpoints. We then tested the classifier on its ability to predict the stimulus identities from neural activity evoked by the third viewpoint in the fourth run of data. Again, we performed a four-fold cross-validation procedure, and also repeated the analysis three times with each viewpoint used as the held out test viewpoint once. We determined the final decoding accuracy by averaging over the four cross-validation iterations and the three viewpoint training and testing combinations.

#### 2.9.3. Identity classification across face and body stimuli

We investigated which regions contain patterns of activity evoked by the stimulus identities that can generalize across neural activity evoked by faces and bodies. We performed these classification analyses separately for neural activity while participants performed the identity and viewpoint recognition tasks. We trained a linear SVM classifier to distinguish between patterns of neural activity evoked by the three face identities using three runs of fMRI data. We then tested the classifier on its ability to predict the identity of the body stimuli in the fourth run of data. We performed a four-fold cross-validation procedure, and also repeated the analysis using neural activity evoked by bodies for training the classifier and neural activity evoked by faces for testing it. We determined the final decoding accuracy by averaging over the four cross-validation iterations and the two training and test set combinations.

## 3. Results

### 3.1. Behavioural results

#### 3.1.1. Identity recognition

Participants’ performance in the identity recognition task was high for both face (96.2 %) and body (93.7 %) stimuli (Fig. 2A, 2C). We investigated if there were any differences in our participants’ ability to recognise the three identities. One-way repeated measures ANOVAs with three levels (ID1, ID2 and ID3) showed that there were no significant differences in participants’ ability to recognise the three identities from the face (*F*_2,36_ = 0.53, *p* = .59, η_p_^2^ = 0.029) or body (*F*_2,36_ = 1.20, *p* = .31, η_p_^2^ = 0.063) stimuli. These results show that participants could easily recognise all stimuli identities from both the faces and bodies.

**Figure 2.**
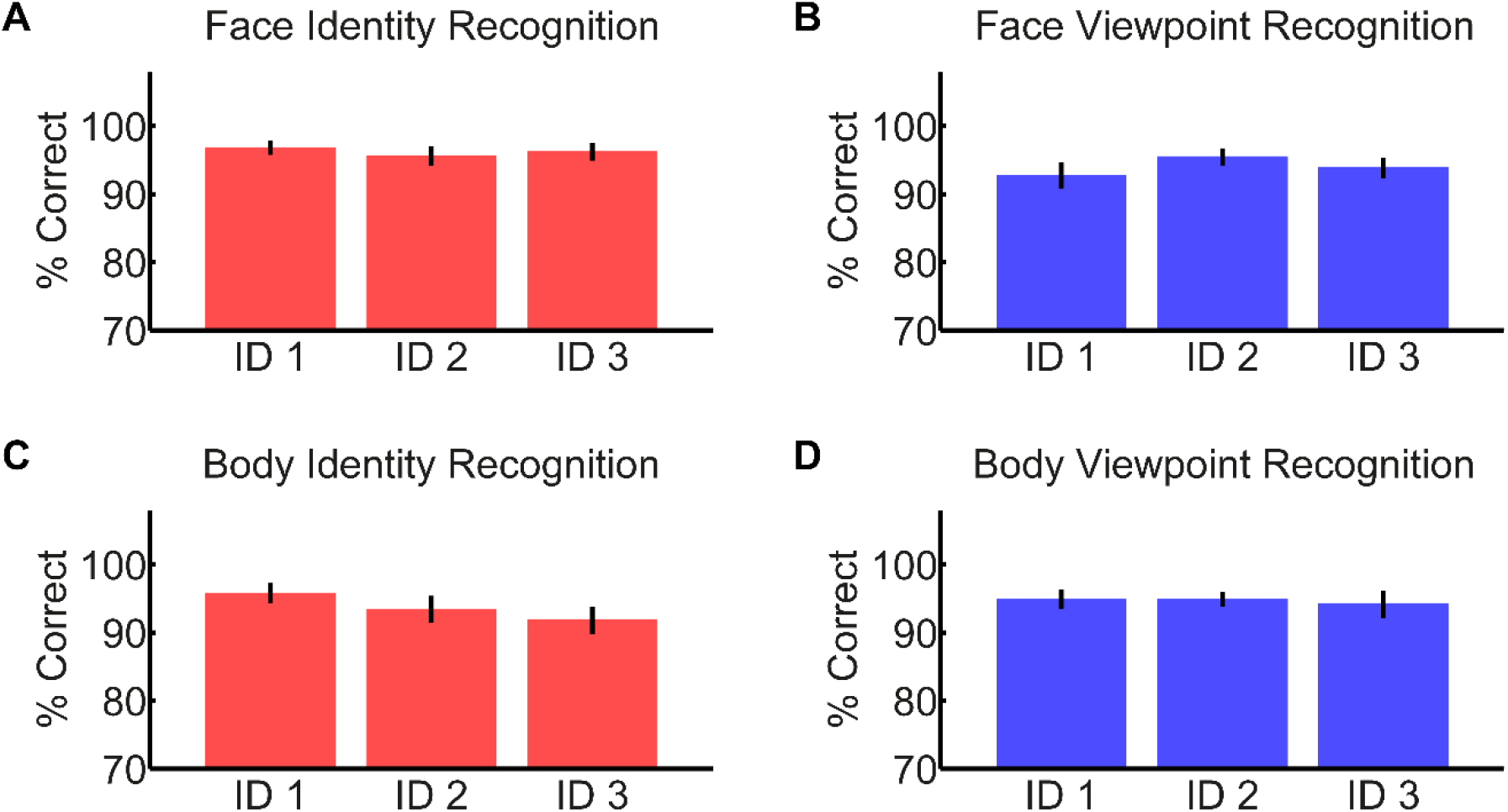
Recognition of the identity and viewpoint of the three stimulus identities (ID1, ID2 & ID3). (A) and (C) show identity recognition accuracy (% correct) for the three stimulus identities from the face (A) and body (C) images. (B) and (D) show viewpoint recognition accuracy (% correct) for the three stimulus identities from the face (B) and body (D) images. Error bars indicate ±1 SEM.

#### 3.1.2. Viewpoint recognition

Participants showed high viewpoint recognition performance for both face (94.0 %) and body (94.6 %) stimuli (Fig. 2B, 2D). One-way repeated measures ANOVAs with three levels (ID1, ID2 and ID3) revealed no significant effect of identity on participants’ ability to recognise the viewpoint of faces (*F*_2,36_ = 2.04, *p* = .14, η_p_^2^ = 0.10) or bodies (*F*_2,36_ = 0.18, *p* = .84, η_p_^2^ = 0.010). Therefore, participants could recognise the stimulus viewpoints equally well regardless of the stimulus identity.

### 3.2. Univariate fMRI results

To investigate whether the three identities evoked different mean levels of BOLD activity, we performed one-way repeated measures ANOVAs with 3 levels (ID1, ID2 and ID3) in face- and body-responsive ROIs and in whole-brain analyses. The results are shown in Figure 3.

**Figure 3.**
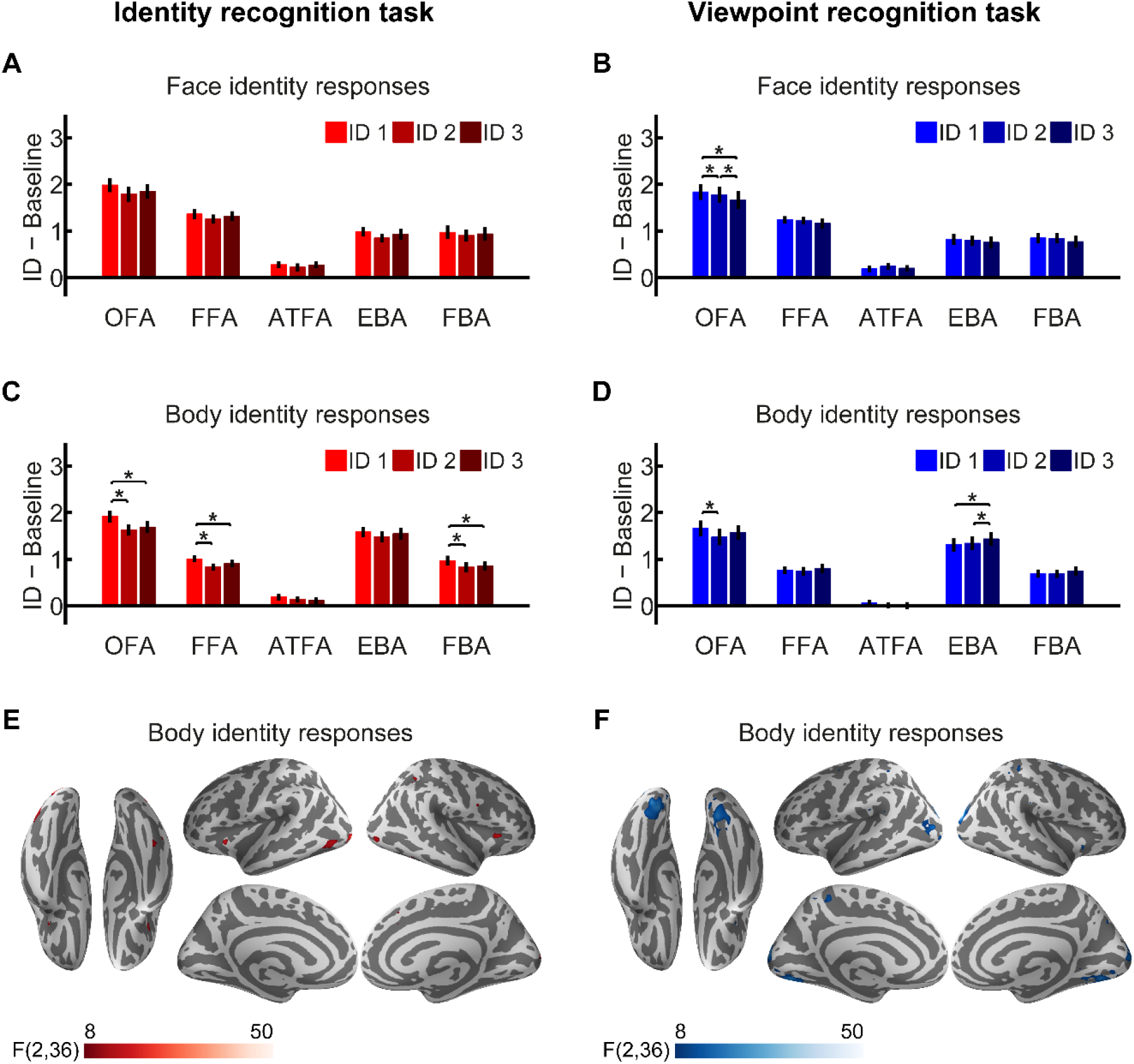
Differences in mean BOLD response to the three identities. (A) and (B) show mean BOLD responses to the three face identities in face- and body-responsive ROIs during the identity (A) and viewpoint (B) recognition task. (C) and (D) show mean BOLD responses to the three body identities in face- and body-responsive ROIs during the identity (C) and viewpoint (D) recognition task. (E) and (F) show differences in mean BOLD responses to the three body identities in whole-brain analyses (FDR corrected) during the identity (E) and viewpoint (F) recognition task. * indicates *p* < .05.

#### 3.2.1. Face identity responses

We performed one-way repeated measures ANOVAs with 3 levels (ID1, ID2 and ID3) to test whether there were any differences in the mean BOLD activity evoked by the three face identities. Full results of these ANOVAs in our ROIs are shown in Table 2. For the identity recognition task, we found no significant differences between the mean BOLD activity evoked by the three face identities in any of our face- or body-responsive ROIs (Fig. 3A) or in any other region in a whole-brain analysis. For the viewpoint recognition task (Fig. 3B), we found significant differences between the mean BOLD activity evoked by the three face identities in the OFA (*F*_2,36_ = 10.27, *p* = .0075) but not in any other ROIs. Follow-up paired *t*-tests showed that in the OFA there was higher activity to ID1 compared to ID2 (*M* = 0.056, *SE* = 0.023, *t_18_* = 2.43, *p* = .026, Cohen’s *d* = 0.56) and ID3 (*M* = 0.17, *SE* = 0.042, *t_18_* = 4.03, *p* < .001, Cohen’s *d* = 0.93), and higher activity to ID2 than ID3 (*M* = 0.11, *SE* = 0.045, *t_18_* = 2.50, *p* = .023, Cohen’s *d* = 0.57). In addition, a whole-brain analysis identified small, bilateral clusters in the early visual cortex showing differences in BOLD activity to the three face identities during the viewpoint recognition task.

**Table 2.** Results of one-way repeated measures ANOVAs testing the effect of face identity on mean BOLD responses in face and body-responsive ROIs.

| Behavioural task | ROI | $F_{2,36}$ | $p$ (corrected) | $p$ (uncorrected) | $\eta_p^2$ |
| --- | --- | --- | --- | --- | --- |
| Identity recognition task | OFA | 6.53 | .056 | .011 | 0.27 |
|  | FFA | 3.52 | .201 | .040 | 0.16 |
|  | ATFA | 1.52 | 1.00 | .234 | 0.08 |
|  | EBA | 4.51 | .089 | .018 | 0.20 |
|  | FBA | 1.71 | .977 | .196 | 0.09 |
| Viewpoint recognition task | OFA | 10.27 | .008 | .002 | 0.36 |
|  | FFA | 2.47 | .594 | .119 | 0.12 |
|  | ATFA | 0.99 | 1.00 | .382 | 0.05 |
|  | EBA | 1.55 | 1.00 | .227 | 0.08 |
|  | FBA | 2.70 | .484 | .097 | 0.13 |

Note. $P$ (corrected) represents $p$ values Bonferroni-corrected for multiple comparisons ( $N = 5$ ROIs). All $p$ values are Greenhouse-Geisser corrected for any cases of non-sphericity.

#### 3.2.2. Body identity responses

We performed the same analysis on the neural responses evoked by body identities. ROI results are shown in Table 3. For the identity recognition task (Fig. 3C), we found significant differences in the mean BOLD responses evoked by the three body identities in the FBA (*F*_2,36_ = 6.96, *p* = .014), OFA (*F*_2,36_ = 20.76, *p* < .001) and FFA (*F*_2,36_ = 11.21, *p* < .001), but not in the EBA or ATFA. Follow-up paired *t*-tests revealed higher activity to ID1 compared to ID2 (FBA: *M* = 0.14, *SE* = 0.028, *t_18_* = 5.03, *p* < .001, Cohen’s *d* = 1.15; OFA: *M* = 0.29, *SE* = 0.044, *t_18_* = 6.56, *p* < .001, Cohen’s *d* = 1.50; FFA: *M* = 0.17, *SE* = 0.030, *t_18_* = 5.67, *p* < .001, Cohen’s *d* = 1.30) and ID3 (FBA: *M* = 0.11, *SE* = 0.047, *t_18_* = 2.40, *p* = .027, Cohen’s *d* = 0.55; OFA: *M* = 0.23, *SE* = 0.046, *t_18_* = 4.98, *p* < .001, Cohen’s *d* = 1.14; FFA: *M* = 0.10, *SE* = 0.038, *t_18_* = 2.67, *p* = .016, Cohen’s *d* = 0.61) but no difference between activity to ID2 and ID3 (FBA: *M* = −0.028, *SE* = 0.043, *t_18_* = −0.66, *p* = .52, Cohen’s *d* = −0.15; OFA: *M* = −0.058, *SE* = 0.051, *t_18_* = −1.13, *p* = .27, Cohen’s *d* = −0.26; FFA: *M* = −0.071, *SE* = 0.041, *t_18_* = −1.74, *p* = .098, Cohen’s *d* = −0.40). We performed a whole-brain analysis to investigate if there were any additional regions showing different levels of mean BOLD activity to the three body identities during the identity recognition task (Fig. 3E). We identified bilateral clusters in the early visual cortex, occipitotemporal cortex (overlapping with the locations of the OFA, FFA and FBA) and insula cortex, and unilateral clusters in the right inferior parietal cortex, right precuneus and right medial superior frontal gyrus.

**Table 3.** Results of one-way repeated measures ANOVAs testing the effect of body identity on mean BOLD responses in face and body-responsive ROIs.

| Behavioural task | ROI | $F_{2,36}$ | $p$ (corrected) | $p$ (uncorrected) | $\eta_p^2$ |
| --- | --- | --- | --- | --- | --- |
| Identity recognition task | OFA | 20.76 | < .001 | < .001 | 0.54 |
|  | FFA | 11.21 | < .001 | < .001 | 0.38 |
|  | ATFA | 1.75 | .942 | .188 | 0.09 |
|  | EBA | 2.02 | .740 | .148 | 0.10 |
|  | FBA | 6.96 | .014 | .003 | 0.28 |
| Viewpoint recognition task | OFA | 6.52 | .019 | .004 | 0.27 |
|  | FFA | 1.43 | 1.00 | .253 | 0.07 |
|  | ATFA | 2.27 | .588 | .118 | 0.11 |
|  | EBA | 6.11 | .026 | .005 | 0.25 |
|  | FBA | 1.88 | .834 | .167 | 0.10 |

Note. $P$ (corrected) represents $p$ values Bonferroni-corrected for multiple comparisons ( $N = 5$ ROIs). All $p$ values are Greenhouse-Geisser corrected for any cases of non-sphericity.

For the viewpoint recognition task (Fig. 3D), we found significant differences in activity evoked by the three body identities in the OFA (*F*_2,36_ = 6.52, *p* = .019) and EBA (*F*_2,36_ = 6.11, *p* = .026), but not in any other ROIs. Follow-up paired *t*-tests showed lower activity in the OFA to ID2 compared to ID1 (*M* = −0.19, *SE* = 0.053, *t_18_* = −3.55, *p* = .0023, Cohen’s *d* = - 0.81) and higher activity in the EBA to ID3 compared to both ID1 (*M* = 0.12, *SE* = 0.036, *t_18_* = 3.43, *p* = .0030, Cohen’s *d* = 0.79) and ID2 (*M* = 0.091, *SE* = 0.036, *t_18_* = 2.51, *p* = .022, Cohen’s *d* = 0.58). We performed a whole-brain analysis to investigate if any other regions would show different levels of mean response to the three body identities during the viewpoint recognition task. We identified bilateral clusters in the early visual cortex, middle occipital cortex, fusiform gyrus, superior temporal cortex, superior parietal cortex, precuneus, superior frontal cortex and insula cortex (Fig. 3F).

### 3.3. Face identity MVPA

To investigate which brain regions contain separable patterns of neural responses for individual face identities, we performed multivoxel pattern analyses in face- and body-responsive ROIs and in whole-brain searchlight analyses. Specifically, we investigated whether face identity could be classified from patterns of neural activity, and whether identity classification could generalise across viewpoints. The results are shown in Figure 4.

**Figure 4.**
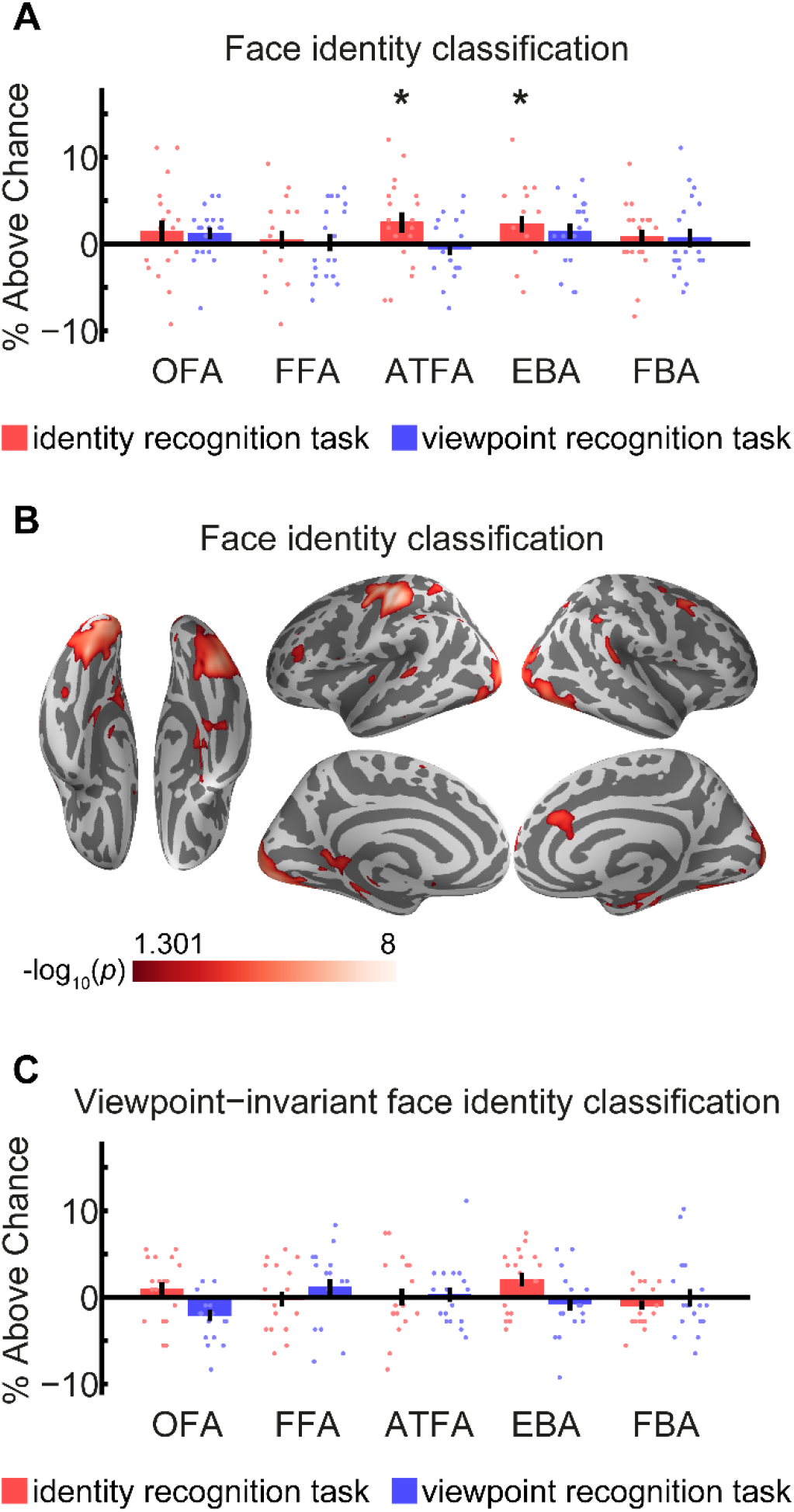
Classification and viewpoint-invariant classification of face identity. (A) shows face identity classification above chance-level (1/3) in face- and body-responsive ROIs. (B) shows classification of face identity during the identity recognition task in a whole-brain searchlight analysis. (C) shows viewpoint-invariant face identity classification above chance-level (1/3) in face- and body-responsive ROIs. Scatter points in (A) and (C) show classification accuracies for individual participants, error bars show ±1 SEM, * indicates *p* < .05 Bonferroni corrected. The colour scale bar in (B) shows −log_10_(*p* values) ranging between 1.301 (*p* = .05) and 8 (*p* = 1 x10^-8^), FDR corrected.

#### 3.3.1. Face identity classification

We first performed ROI-based MVPA to investigate which face- and body responsive ROIs could decode face identity above chance-level (i.e. 1/3; see Section 2.9.1 for details of methods). The results are shown in Fig. 4A (see Table 4 for full results). From the identity recognition task data, we were able to decode face identity significantly above chance from the face-responsive ATFA (35.8 %, *p* = .031) and body-responsive EBA (35.6 %, *p* = .045), but not from other face-responsive ROIs or the FBA. From the viewpoint recognition task data, we were not able to decode face identity from any ROIs.

**Table 4.** Classification of face identity from patterns of neural responses in individual ROIs.

| Analysis | Behavioural task | ROI | Classification accuracy (%) | $p$ (corr.) | $p$ (uncorr.) | Cohen's $d$ |
| --- | --- | --- | --- | --- | --- | --- |
| Face identity classification | Identity recognition task | OFA | 34.8 | .318 | .064 | 0.27 |
|  |  | FFA | 33.8 | 1.00 | .309 | 0.11 |
|  |  | ATFA | 35.8 | .031 | .006 | 0.49 |
|  |  | EBA | 35.6 | .045 | .009 | 0.56 |
|  |  | FBA | 34.1 | 1.00 | .201 | 0.20 |
|  | Viewpoint recognition task | OFA | 34.6 | .505 | .101 | 0.40 |
|  |  | FFA | 33.5 | 1.00 | .443 | 0.03 |
|  |  | ATFA | 32.8 | 1.00 | .705 | -0.16 |
|  |  | EBA | 34.8 | .335 | .067 | 0.37 |
|  |  | FBA | 34.0 | 1.00 | .224 | 0.15 |
| Viewpoint-invariant face identity classification | Identity recognition task | OFA | 34.3 | .826 | .165 | 0.26 |
|  |  | FFA | 33.1 | 1.00 | .617 | -0.07 |
|  |  | ATFA | 33.4 | 1.00 | .488 | 0.01 |
|  |  | EBA | 35.4 | .059 | .012 | 0.60 |
|  |  | FBA | 32.4 | 1.00 | .862 | -0.40 |
|  | Viewpoint recognition task | OFA | 31.2 | 1.00 | .991 | -0.77 |
|  |  | FFA | 34.5 | .551 | .110 | 0.27 |
|  |  | ATFA | 33.6 | 1.00 | .373 | 0.08 |
|  |  | EBA | 32.7 | 1.00 | .773 | -0.19 |
|  |  | FBA | 33.3 | 1.00 | .513 | -0.01 |

*Note.* Statistical significance was assessed using permutation tests, and $p$ values are shown before (uncorrected) and after (corrected) Bonferroni-correction for multiple comparisons ( $N = 5$ ROIs).

Secondly, we performed a whole-brain searchlight analysis to investigate if any other brain regions could decode face identity. Using fMRI data from the identity recognition task we identified clusters than could decode face identity bilaterally in the early visual cortex, inferior occipital cortex, fusiform gyrus, superior parietal cortex, superior temporal cortex and parahippocampal gyrus, and unilaterally in the right middle frontal gyrus, right anterior cingulum, right medial superior frontal gyrus and left inferior frontal gyrus (Fig. 4B). We could also decode identity in the left motor cortex as participants pressed different buttons for each stimulus identity. Using fMRI data from the viewpoint recognition task, we were unable to decode face identity from any regions.

#### 3.3.2. Viewpoint-invariant face identity classification

We next investigated which regions could classify face identity across viewpoint using the methods described in Section 2.9.2. Results of this analysis in face- and body-responsive ROIs are summarised in Table 4. We were unable to decode face identity across viewpoint from any of the ROIs we tested (Fig. 4C) using fMRI data from either the identity recognition task or the viewpoint recognition task. We performed searchlight analyses to investigate if any other brain regions would be able to decode face identity across viewpoint. We did not identify any regions in these analyses.

### 3.4. Body identity MVPA

We performed the same MVPA analyses for bodies as for faces, to investigate which brain regions contain separable patterns of neural responses evoked by different body identities, and to investigate whether the neural coding of body identity can generalize across different viewpoints. We performed these analyses in face- and body-responsive ROIs and in whole-brain searchlight analyses.

#### 3.4.1. Body identity classification

First, we investigated which of our face- and body-responsive ROIs could decode body identity above chance-level (1/3; see Section 2.9.1 for method details). The results are shown in Fig. 5A (see Table 5 for full results of ROI-based classification of body identity). From the identity recognition task, we could decode body identity significantly above chance from the body-responsive FBA (36.4 %, *p* = .0045) and face-responsive OFA (38.5 %, *p* < .001), but not from the body-responsive EBA or any other ROIs. From the viewpoint recognition task, we were able to decode body identity from the OFA (40.7 %, *p* < .001), but not from any other ROIs.

**Figure 5.**
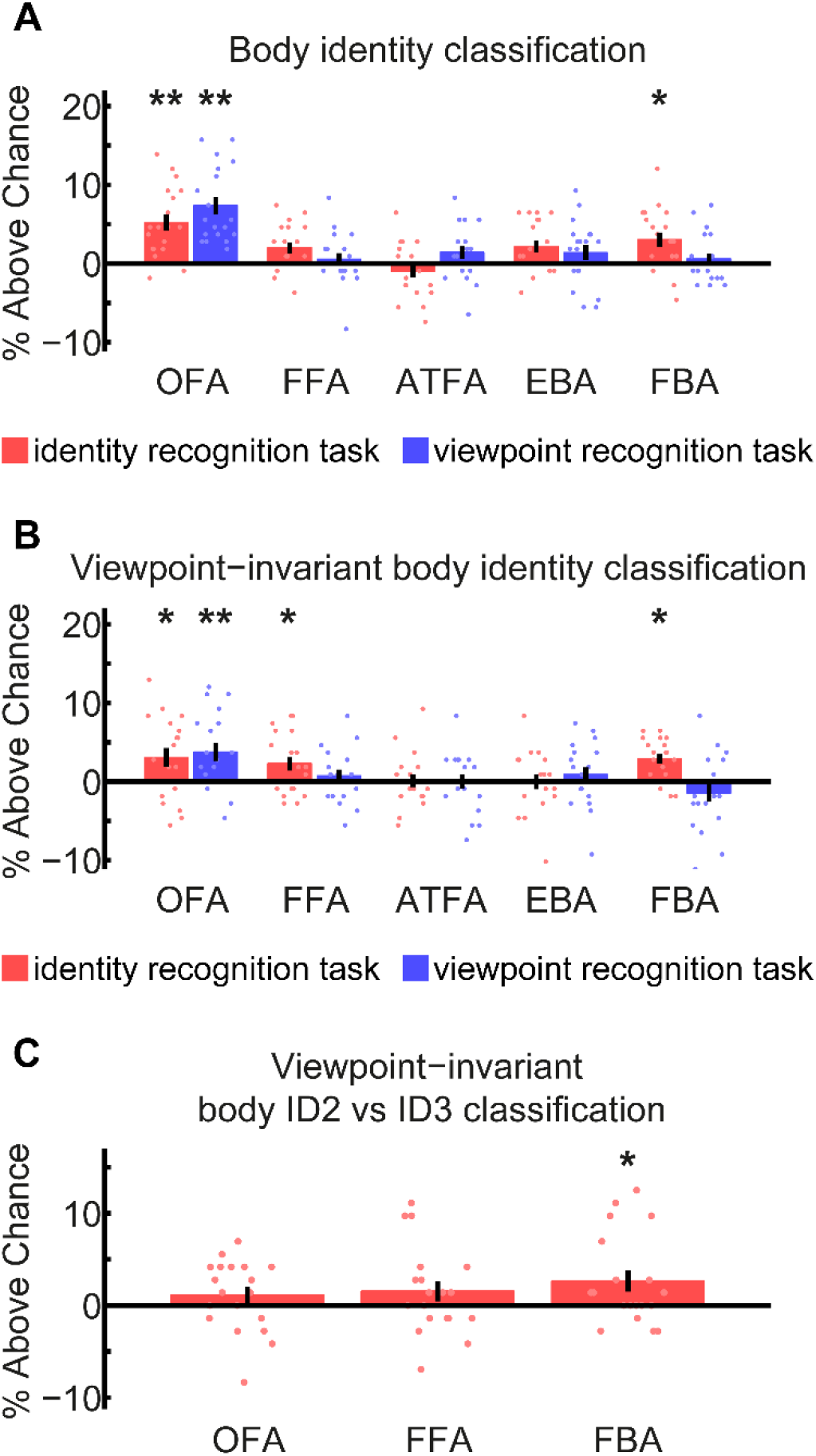
Classification and viewpoint-invariant classification of body identity in face- and body-responsive ROIs. (A) shows body identity classification above chance-level (1/3), (B) shows viewpoint-invariant body identity classification above chance-level (1/3) and (C) shows viewpoint-invariant body identity classification above chance-level (1/2) for ID2 vs. ID3 only, with fMRI data from the identity recognition task. Scatter points show classification accuracies for individual participants and error bars show ±1 SEM. ** indicates *p* < .001, * indicates *p* < .05, Bonferroni corrected.

**Table 5.** Classification of body identity from patterns of neural responses in individual ROIs.

| Analysis | Behavioural task | ROI | Classification accuracy (%) | $p$ (corr.) | $p$ (uncorr.) | Cohen's $d$ |
| --- | --- | --- | --- | --- | --- | --- |
| Body identity classification | Identity recognition task | OFA | 38.5 | < .001 | < .001 | 1.18 |
|  |  | FFA | 35.3 | .091 | .018 | 0.67 |
|  |  | ATFA | 32.4 | 1.00 | .842 | -0.28 |
|  |  | EBA | 35.5 | .071 | .014 | 0.65 |
|  |  | FBA | 36.4 | .005 | < .001 | 0.76 |
|  | Viewpoint recognition task | OFA | 40.7 | < .001 | < .001 | 1.54 |
|  |  | FFA | 33.9 | 1.00 | .282 | 0.16 |
|  |  | ATFA | 34.7 | .347 | .069 | 0.39 |
|  |  | EBA | 34.7 | .390 | .078 | 0.33 |
|  |  | FBA | 33.9 | 1.00 | .273 | 0.19 |
| Viewpoint-invariant body identity classification | Identity recognition task | OFA | 36.4 | .004 | < .001 | 0.59 |
|  |  | FFA | 35.6 | .024 | .005 | 0.60 |
|  |  | ATFA | 33.4 | 1.00 | .468 | 0.03 |
|  |  | EBA | 33.3 | 1.00 | .535 | 0.01 |
|  |  | FBA | 36.2 | .003 | < .001 | 1.06 |
|  | Viewpoint recognition task | OFA | 37.1 | < .001 | < .001 | 0.76 |
|  |  | FFA | 34.1 | 1.00 | .215 | 0.21 |
|  |  | ATFA | 33.3 | 1.00 | .499 | 0.00 |
|  |  | EBA | 34.2 | .783 | .157 | 0.22 |
|  |  | FBA | 31.9 | 1.00 | .955 | -0.30 |

*Note.* Statistical significance was assessed using permutation tests, and $p$ values are shown before (uncorrected) and after (corrected) Bonferroni-correction for multiple comparisons ( $N = 5$ ROIs).

Second, we performed a whole-brain searchlight analysis to investigate if we could decode body identity from any other brain regions. As illustrated in Fig. 6A, we could decode body identity from a large area of occipital cortex using fMRI data from both the identity and viewpoint recognition tasks. Using fMRI data from the identity recognition task, we could also decode body identity from bilateral regions in the fusiform gyrus, superior parietal cortex, inferior frontal gyrus and middle frontal gyrus, and unilaterally from the right anterior temporal cortex and right insula cortex. We could also decode body identity in the left motor cortex as participants pressed different buttons to indicate the stimulus identity. Using fMRI data from the viewpoint recognition task, we could also decode body identity from bilateral regions in the fusiform gyrus, superior parietal cortex, supramarginal gyrus, cingulum, precentral gyrus and the caudate nucleus, and unilaterally from the right superior frontal gyrus.

**Figure 6.**
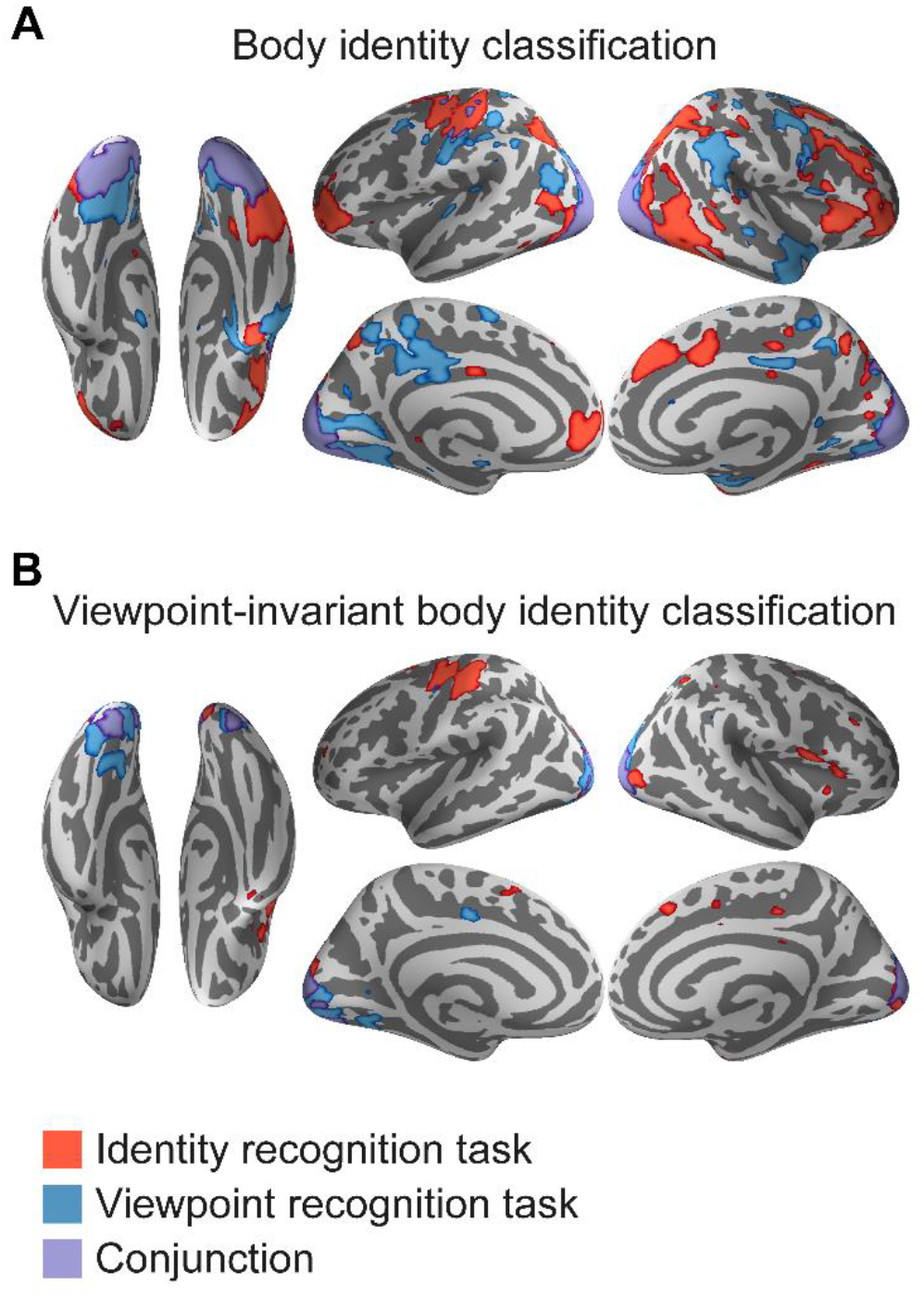
Classification (A) and viewpoint-invariant classification (B) of body identity in whole-brain searchlight analyses. Regions showing significant activity during the identity recognition task are shown in red, regions showing significant activity during the viewpoint recognition task are shown in blue and regions showing significant activity during both tasks (conjunction) are shown in purple. Significant regions were defined using a *p* < .05 FDR correction.

#### 3.4.2. Viewpoint-invariant body identity classification

We first tested which of our face- and body-responsive ROIs could decode body identity across viewpoint (Fig. 5B & Table 5; for details of methods, see Section 2.9.2). From the identity recognition task, we could decode body identity across viewpoint significantly above chance-level (1/3) in the body-responsive FBA (36.2 %, *p* = .0030) and face-responsive OFA (36.4 %, *p* = .0035) and FFA (35.6 %, *p* = .024). We were not able to decode body identity across viewpoint from the body-responsive EBA or the face-responsive ATFA. From the viewpoint recognition task, we could decode body identity across viewpoint from the OFA (37.1 %, *p* < .001), but not from any other ROI.

We performed a whole-brain searchlight analysis to investigate if any other brain regions could decode body identity across viewpoint (Fig. 6B). From both the identity and the viewpoint task data, we could decode body identity across viewpoint from a large cluster in occipital cortex (including the early visual cortex). Using fMRI data from the identity recognition task, we could additionally decode body identity across viewpoint from the middle frontal gyrus, right anterior temporal cortex, right superior parietal cortex, right medial superior frontal gyrus, right insula cortex, right rolandic operculum and the left motor cortex (due to participants’ button presses). Using fMRI data from the viewpoint recognition task, we could additionally decode body identity across viewpoint from the left fusiform gyrus, right superior parietal cortex, left caudate nucleus, left cingulum and left postcentral gyrus.

#### 3.4.3. Viewpoint-invariant body identity classification: ID2 vs. ID3

As we found higher BOLD responses to ID1 as compared to ID2 and ID3 in the OFA, FFA and FBA during the identity response task (Fig. 3C), it is possible that our body identity decoding across viewpoint results in these regions was driven by differences in univariate activation. To explore whether purely multivariate pattern differences also contributed to body identity decoding across viewpoint in these regions, we performed an additional analysis to test if these regions would be able to classify only body identities ID2 and ID3 across viewpoint (Fig. 5C). We performed the analysis using the same method as in *Section 3.4.2*, except that we only trained and tested the classifiers ability to distinguish between ID2 and ID3. We were able to decode body identity across viewpoint significantly above chance (1/2) in the body-responsive FBA (52.6 %, *p* = .038 Bonferroni corrected, Cohen’s *d* = 0.54) but not in the face-responsive OFA (51.1 %, *p* = .17 uncorrected, Cohen’s *d* = 0.28) or FFA (51.5 %, 0.097 uncorrected, Cohen’s *d* = 0.32).

### 3.5. Identity classification across face and body stimuli

Lastly, we performed multivoxel pattern analyses to investigate if any brain regions contain patterns of neural activity evoked by the identity of a person that could generalize across neural activity evoked by face and body stimuli (see Section 2.9.3 for method details). We performed the analyses in face- and body-responsive ROIs (Fig. 7A & Table 6) and in whole-brain searchlight analyses (Fig. 7B).

**Figure 7.**
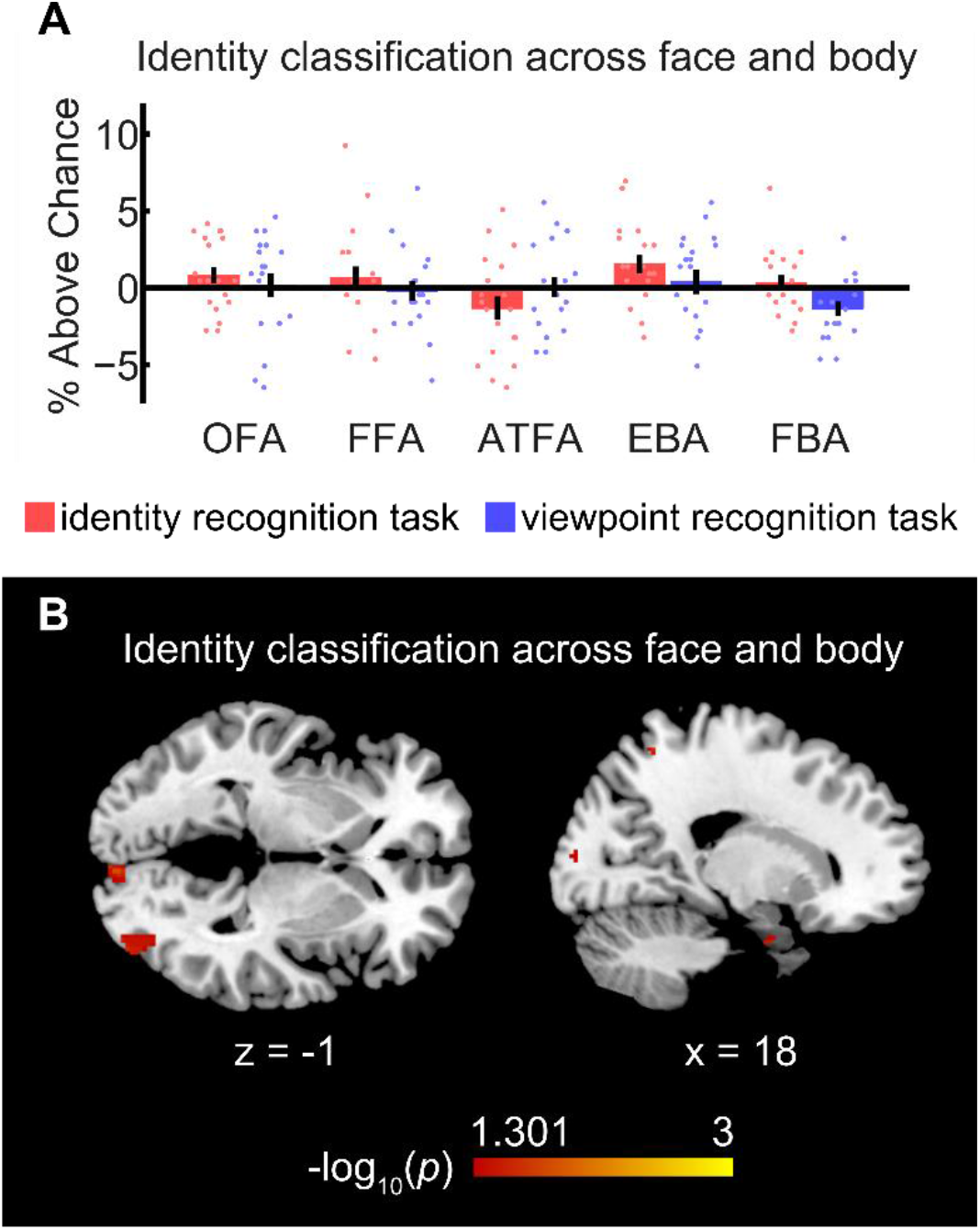
Classification of identity across neural activity evoked by faces and bodies. (A) shows classification of identity above chance-level (1/3) across face and body stimuli in face- and body-responsive ROIs. Scatter points show classification accuracies for individual participants and error bars show ±1 SEM. (B) shows classification of identity across face and body stimuli during the identity recognition task in a whole-brain searchlight analysis. The scale bar shows −log_10_(*p* values) between 1.301 (*p* = .05) and 3 (*p* = .001), FDR corrected.

**Table 6.** Classification of identity across neural activity evoked by face and body stimuli in ROIs.

| Behavioural task | ROI | Classification accuracy (%) | $p$ (corr.) | $p$ (uncorr.) | Cohen's $d$ |
| --- | --- | --- | --- | --- | --- |
| Identity recognition task | OFA | 34.2 | .917 | .183 | 0.35 |
|  | FFA | 34.0 | 1.00 | .231 | 0.20 |
|  | ATFA | 32.0 | 1.00 | .926 | -0.40 |
|  | EBA | 34.9 | .210 | .042 | 0.58 |
|  | FBA | 33.7 | 1.00 | .339 | 0.16 |
| Viewpoint recognition task | OFA | 33.5 | 1.00 | .425 | 0.05 |
|  | FFA | 33.1 | 1.00 | .579 | -0.07 |
|  | ATFA | 33.4 | 1.00 | .484 | 0.01 |
|  | EBA | 33.7 | 1.00 | .332 | 0.11 |
|  | FBA | 32.0 | 1.00 | .935 | -0.67 |

*Note.* Statistical significance was assessed using permutation tests, and $p$ values are shown before (uncorrected) and after (corrected) Bonferroni-correction for multiple comparisons ( $N = 5$ ROIs).

Irrespective of using fMRI data from the identity recognition task or the viewpoint recognition task, we were unable to decode identity across neural activity evoked by faces and bodies higher than chance-level (1/3) in any of the ROIs we tested (Fig. 7A & Table 6). We performed whole-brain searchlight analyses to investigate if any other brain regions could decode identity across neural activity evoked by faces and bodies. Using fMRI data from the identity recognition task (Fig. 7B), we could decode identity from the early visual cortex (MNI: 10, −94, 2), a region in the right inferior occipital cortex (MNI: 40, −84, −4) overlapping with the mean location of the OFA, the right parahippocampal cortex (MNI: 20, −4, −30) and a region in the right superior parietal cortex (MNI: 16, −56, 60). We could also decode identity from the left motor cortex due to participants’ button presses. In contrast, when using fMRI data from the viewpoint recognition task, we were not able to decode identity from any regions.

## 4. Discussion

In this study, we investigated how face and body identities are encoded in the brain, and whether person identity is encoded in an abstract neural representation that generalises across face and body stimuli. Consistent with previous findings, we found that face identity could be decoded from neural activity in several distributed cortical regions (Anzellotti & Caramazza, 2016). We found that body identity could also be decoded from neural activity in several distributed cortical regions, and in particular we found consistent decoding of body identity, including across viewpoint, from neural activity in the FBA, the right anterior temporal cortex, the middle frontal gyrus and the right insula cortex. We found we could decode identity in an abstract manner, across neural activity evoked by faces and bodies, from neural activity in the right parahippocampal cortex, right superior parietal cortex, right inferior occipital cortex and early visual cortex. These results provide new insights into how the brain encodes information about person identity.

### 4.1. Neural coding of face identity

Our ROI-based analysis showed that face identity can be decoded from neural activity in the face-responsive ATFA and body-responsive EBA. Our whole-brain searchlight analysis revealed a broader and more distributed brain network involved in encoding face identity, including the early visual cortex, inferior occipital cortex, fusiform gyrus, superior parietal cortex, superior temporal cortex, parahippocampal cortex, right middle frontal gyrus, right anterior cingulum, right medial superior frontal gyrus and left inferior frontal gyrus. These results are consistent with previous findings, which have shown that face identity can be decoded from a number of distributed brain regions, including the ATFA, FFA, OFA, superior intraparietal sulcus and right inferior frontal cortex (Anzellotti & Caramazza, 2016; Anzellotti et al., 2014; Axelrod & Yovel, 2015; Goesaert & Op de Beeck, 2013; Guntupalli et al., 2017; Jeong & Xu, 2016; Kriegeskorte et al., 2007; Natu et al., 2010; Nestor et al., 2011). Note that we could decode face identity only when participants attended to the identity of the stimuli, but not when they attended to the stimulus viewpoint. This suggests that face identity decoding in these regions was not due to differences in visual features, as these were identical in both tasks (see Section 4.4 for a discussion of the behavioural task differences).

Neither our ROI analysis nor our searchlight analysis showed above-chance decoding of face identity across viewpoint. Electrophysiological recordings in macaque monkeys have shown that neurons in the anterior face patches respond to face identity across viewpoint (Freiwald & Tsao, 2010). Correspondingly, human neuroimaging studies have shown that face identity can be decoded across viewpoint from human face-responsive regions (Anzellotti et al., 2014; Guntupalli et al., 2017). One possibility for the discrepancy between these findings and our results is that people may need more extensive learning to develop viewpoint-invariant coding of face identity. In comparison to a training session of 30 minutes (Anzellotti et al., 2014) or with 360 trials (Guntupalli et al., 2016), our training (30 minutes for both face and body learning, with 135 face trials in total) might be insufficient for participants to establish a viewpoint-independent neural representation of face identity (despite high behavioural performance). Previous work has also demonstrated that in some cases fMRI MVPA can fail to decode identity, even when electrophysiological recordings show that viewpoint-invariant identity information is present in the underlying neurons (Dubois, de Berker, & Tsao, 2015).

### 4.2. Neural coding of body identity

We could decode body identity from the body-responsive FBA and the face-responsive OFA and FFA in our ROI analyses, and from regions in the occipital cortex, fusiform gyrus, right anterior temporal cortex, superior parietal cortex, supramarginal gyrus, cingulum, precentral gyrus, caudate nucleus, inferior frontal gyrus, middle frontal gyrus, right insula cortex and right superior frontal gyrus in our searchlight analyses. Several of these regions have been shown to have higher responses to bodies of familiar people as compared to unfamiliar people (Hodzic et al., 2009). We could decode body identity from the OFA and occipital cortex regardless of the recognition task, suggesting these regions may encode body identities using visual body features, and that their neural coding might not be influenced by top down factors such as attention. In contrast, decoding of body identity from the right anterior temporal cortex, inferior frontal gyrus, middle frontal gyrus and right insula was only possible when participants attended to identity, suggesting that body identity coding in these regions was enhanced by participants’ attention to identity, rather than being driven solely by visual features.

In contrast to the decoding of face identity, our ROI analyses revealed viewpoint-tolerant encoding of body identity from neural activity in the FBA, OFA and FFA. Furthermore, viewpoint-tolerant coding of body identity was evident in more distributed regions in our searchlight analyses, including the occipital cortex, middle frontal gyrus, right anterior temporal cortex, right superior parietal cortex, right medial superior frontal gyrus, right insula cortex, right rolandic operculum, left caudate nucleus, left cingulum and left postcentral gyrus. Remarkably, the OFA and occipital cortex showed consistent above-chance decoding of body identity across viewpoints regardless of recognition tasks, suggesting that these regions may encode body identity using visual features that can generalise across different viewpoints. In contrast, attention to body identity was required to decode body identity across viewpoint from the FBA, FFA, right anterior temporal cortex, middle frontal gyrus, right medial superior frontal gyrus, right insula cortex and right rolandic operculum, suggesting that attention to body identity may have enhanced its neural coding in these regions.

Among the brain regions showing above chance decoding of body identities, FBA, FFA, OFA, right medial superior frontal gyrus and right insula also showed different univariate responses to the three body identities. For our ROIs, we performed a further analysis to disentangle whether the decoding of body identity in these regions was driven by purely univariate responses or also by multivariate differences in the pattern of neural responses. We found that we could decode body identity across viewpoint in the FBA, but not the FFA or OFA, between two identities that showed no difference in their univariate responses, demonstrating that multivariate patterns also contribute to body identity decoding across viewpoint in the FBA. Altogether, these results demonstrate a robust viewpoint-invariant neural encoding of body identity in the FBA, the right anterior temporal cortex, the middle frontal gyrus and the right insula cortex. All these regions showed consistent decoding of body identity in two analyses (body identity decoding and viewpoint-invariant body identity decoding), and this decoding was not driven by low-level visual features.

Our finding that body identity can be decoded across viewpoint in the FBA and right anterior temporal cortex is consistent with the view that identity coding in the anterior temporal cortex is tolerant to viewpoint changes. Several previous studies have found viewpoint-invariant responses to face identity in the anterior temporal cortex (Anzellotti et al., 2014; Freiwald & Tsao, 2010; Guntupalli et al., 2017). Similarly, classification of body identity across viewpoint and pose was higher in a more anterior body patch in macaque temporal cortex than a more posterior body patch (Kumar et al., 2019). Our results are consistent with these findings, suggesting that disentangling identity from viewpoint may be a general function of anterior temporal regions. Furthermore, our results also show a functional dissociation between neural coding of body identity in the FBA and the EBA, as we were able to decode body identity in the FBA, but not in the EBA in any of our analyses. Although a previous study found responses to body identity in both EBA and FBA using a repetition-suppression paradigm, feedback connectivity from the FBA to the EBA may have driven the body identity repetition suppression in the EBA in this study (Ewbank et al., 2011). Therefore, correspondingly to the coding of face identity along the posterior-anterior axis, encoding of body identity may also develop from viewpoint-sensitive in the EBA to viewpoint-invariant in the FBA.

### 4.3. Neural coding of identity across face and body

We could decode identity across neural activity evoked by face and body stimuli in the early visual cortex, the right inferior occipital cortex, the right parahippocampal cortex and the right superior parietal cortex. This abstract identity decoding was possible when participants attended to stimulus identity, but not when they attended to viewpoint, showing that this decoding was enhanced by participants’ attention to identity, and was not solely based on visual features. Our ability to decode identity across the face and body in the early visual cortex may be due to feedback of identity information from high-level brain regions to the early visual cortex. Previous studies have demonstrated such feedback of high-level visual information to the early visual cortex (Bannert & Bartels, 2013; Grassi, Zaretskaya, & Bartels, 2017; Zaretskaya, Anstis, & Bartels, 2013). Although we could not decode identity from our OFA ROI, we could decode identity from a region in the right inferior occipital cortex overlapping with the mean location of the right OFA. Together with the finding of viewpoint-invariant coding of face identity in the OFA (Anzellotti et al., 2014), this result suggests that inferior occipital cortex may contain some abstract encoding of person identity. We could also decode identity in an abstract manner in the right parahippocampal cortex, a region is known to be involved in memory and recollection (Eichenbaum, Yonelinas, & Ranganath, 2007). Previous studies have shown an involvement of parahippocampal cortex in identity coding. Famous faces elicit stronger parahippocampal cortex activation as compared to unfamiliar faces (Bar, Aminoff, & Ishai, 2008), and this region is also activated by recollection of contextual associations of faces and names (Kirwan & Stark, 2004). Our results suggest that this region also integrates identity information from the face and body. Finally, the right superior parietal cortex also showed above-chance identity decoding across faces and bodies. Previous work has shown there is abstract coding of face and car identity in the parietal cortex (Jeong & Xu, 2016). In combination, our identity decoding results across faces and bodies revealed a network of brain regions that respond to person identity in an abstract manner. These regions fall mostly outside of the standard face and body-responsive brain regions, suggesting that the integration of face and body identity occurs primarily outside of stimuli-selective brain regions.

### 4.4. Effect of attention on neural coding of person identity

We found that we could more frequently decode identity when participants attended to identity (performed the identity recognition task) than when they did not (performed the viewpoint recognition task). We could decode face identity and decode identity across face and body stimuli when participants attended to identity, but not when they attended to viewpoint. Similarly, we could decode body identity from the FBA, right anterior temporal cortex and middle frontal gyrus when participants attended to identity, but not when they attended to viewpoint. In previous research, studies reporting successful decoding of face identity often used tasks where participants attended to identity (Anzellotti & Caramazza, 2016; Anzellotti et al., 2014; Guntupalli et al., 2017; Jeong & Xu, 2016; Nestor et al., 2011), whereas studies reporting unsuccessful decoding of face identity often used tasks that were unrelated to face recognition (Dubois et al., 2015; Ramírez, Cichy, Allefeld, & Haynes, 2014). Recent studies have demonstrated that attention to face identity enhances neural responses to face identity (Dobs, Schultz, Bülthoff, & Gardner, 2018; Gratton, Sreenivasan, Silver, & D’Esposito, 2013). Our results, in combination with these previous studies, suggest that neural representations of face and body identity are enhanced by attention to identity, perhaps due to activation of identity-responsive neurons, and that this enhancement may be necessary to be able to decode person identity based on patterns of neural responses.

## 5. Conclusion

We show, for the first time to our knowledge, that body identity can be decoded across viewpoint from neural activity in the body-responsive FBA, the right anterior temporal cortex, the middle frontal gyrus and the right insula cortex using MVPA. This result provides evidence that viewpoint-invariant identity coding may be a general function of more anterior regions of the human temporal cortex. Furthermore, we show that identity can be decoded in an abstract manner across neural activity evoked by faces and bodies in several brain regions previously associated with abstract identity coding. These results reveal how face and body identities are encoded, how this neural coding can generalise across viewpoints and how it is modulated by attention. Moreover, our results also shed light upon the neural substrates underlying the development of an abstract person identity representation.

## Acknowledgements

This research was supported by the Max Planck Society, Germany.

## Notes

**Conflict of Interest Statement**: None

### Competing Interest Statement

The authors have declared no competing interest.

